# Resolving the Full Spectrum of Human Genome Variation using Linked-Reads

**DOI:** 10.1101/230946

**Authors:** Patrick Marks, Sarah Garcia, Alvaro Martinez Barrio, Kamila Belhocine, Jorge Bernate, Rajiv Bharadwaj, Keith Bjornson, Claudia Catalanotti, Josh Delaney, Adrian Fehr, Ian T. Fiddes, Brendan Galvin, Haynes Heaton, Jill Herschleb, Christopher Hindson, Esty Holt, Cassandra B. Jabara, Susanna Jett, Nikka Keivanfar, Sofia Kyriazopoulou-Panagiotopoulou, Monkol Lek, Bill Lin, Adam Lowe, Shazia Mahamdallie, Shamoni Maheshwari, Tony Makarewicz, Jamie Marshall, Francesca Meschi, Chris O’keefe, Heather Ordonez, Pranav Patel, Andrew Price, Ariel Royall, Elise Ruark, Sheila Seal, Michael Schnall-Levin, Preyas Shah, Stephen Williams, Indira Wu, Andrew Wei Xu, Nazneen Rahman, Daniel MacArthur, Deanna M. Church

## Abstract

Large-scale population based analyses coupled with advances in technology have demonstrated that the human genome is more diverse than originally thought. To date, this diversity has largely been uncovered using short read whole genome sequencing. However, standard short-read approaches, used primarily due to accuracy, throughput and costs, fail to give a complete picture of a genome. They struggle to identify large, balanced structural events, cannot access repetitive regions of the genome and fail to resolve the human genome into its two haplotypes. Here we describe an approach that retains long range information while harnessing the advantages of short reads. Starting from only ∼1ng of DNA, we produce barcoded short read libraries. The use of novel informatic approaches allows for the barcoded short reads to be associated with the long molecules of origin producing a novel datatype known as ‘Linked-Reads’. This approach allows for simultaneous detection of small and large variants from a single Linked-Read library. We have previously demonstrated the utility of whole genome Linked-Reads (lrWGS) for performing diploid, *de novo* assembly of individual genomes (Weisenfeld et al. 2017). In this manuscript, we show the advantages of Linked-Reads over standard short read approaches for reference based analysis. We demonstrate the ability of Linked-Reads to reconstruct megabase scale haplotypes and to recover parts of the genome that are typically inaccessible to short reads, including phenotypically important genes such as *STRC, SMN*_*1*_ and *SMN*_*2*_. We demonstrate the ability of both lrWGS and Linked-Read Whole Exome Sequencing (lrWES) to identify complex structural variations, including balanced events, single exon deletions, and single exon duplications. The data presented here show that Linked-Reads provide a scalable approach for comprehensive genome analysis that is not possible using short reads alone.

## Introduction

Since the completion of the human genome project, many large scale consortia studies have applied whole genome sequencing to thousands of individuals from diverse populations across the globe, reshaping our understanding of human variation (Auton et al. 2015; Lek et al. 2016; Sudmant et al. 2015). To date, most genome analyses were performed with accurate, high-throughput short reads leading to robust analysis of small variants over non-repetitive parts of the genome, but only providing a small window into the landscape of larger structural variants (SVs). The application of recent technical advances in both sequencing and mapping to genome analysis have revealed that despite extensive information garnered from large population surveys utilizing short read whole genome sequencing (srWGS), we are still under-representing the amount of structural variation in the human population in these short read driven studies (Chaisson et al. 2014, 2017; Huddleston and Eichler 2016; Collins et al. 2017).

The reconstruction of long range haplotypes (phasing) can be important for many biological studies. When analyzing data from rare disease cohorts, knowing if potentially pathogenic variants are in *cis* or *trans* is necessary for interpreting the impact of these variants. Additionally, haplotype information is necessary for understanding allele specific impacts on gene expression (Ramaker et al. 2017). In addition to the value that haplotype information can bring to interpreting variation data, studies also show that this information can be critical for variant identification, particularly for SVs that are heterozygous in a sample (Huddleston and Eichler 2016). The ability to routinely obtain long range haplotype information could be beneficial to genome studies.

The limitations of short reads suggest the need for improved methods for genome analysis. Several long molecule sequencing and mapping approaches have been developed to address these issues (Carneiro et al. 2012; Nakano et al. 2017; Genomics 2017). While they provide powerful data for better understanding genome structure, their high input requirements, error rates and costs make them inaccessible to many applications, particularly those requiring thousands of samples (Chaisson et al. 2017). To address this need, we developed a technology that retains long range information while maintaining the benefits of short read sequencing. The core datatype, Linked-Reads, is generated by performing haplotype limiting dilution of long DNA molecules into >1 million barcoded partitions, synthesizing barcoded sequence libraries within those partitions, and then performing standard short read sequencing in bulk. The limited amount of DNA put into the system, coupled with novel algorithms, allow short reads to be associated with their long molecule of origin, in most cases, with high probability.

The Linked-Read datatype was originally described in (Zheng et al. 2016) using the GemCode™ System. Here we describe improvements over GemCode using the Chromium™ System. These improvements come from increasing the number of barcodes (737,000 to 4 million), and the number of partitions (100,000 to 1 million) as well as improving the biochemistry to substantially reduce coverage bias. These improvements eliminate the need for an additional short-read library. We also describe improvements to our analytical pipeline, Long Ranger™.

We compare reference based analysis on multiple standard control samples using either a single Chromium Linked-Read library or a standard short read library for both whole genome (WGS) and whole exome sequencing (WES) approaches. We demonstrate the ability to construct accurate, multi-megabase haplotypes by coupling long molecule information with heterozygous variants within the sample. We show that a single Chromium library has comparable small variant sensitivity and specificity to standard short read libraries and helps expand the amount of the genome that can be accessed and analyzed. We demonstrate the ability to identify large scale SVs, in control and validation samples, by taking advantage of the long range information provided by the barcoded library. Lastly, we assess the ability to identify variants in archival samples that had been previously assessed by orthogonal methods. These data show that a Chromium Linked-Reads provide more genome information than short reads alone.

## Results

Here we describe both the biochemistry improvements that generate barcoded reads, as well as algorithmic improvements that take advantage of these barcodes. It is important to note that Linked-Reads are paired-end short reads with a barcode on read 1 and can be used by many common short read tools. To fully realize the potential of Linked-Reads, additional algorithms that take advantage of these bar coded sequences and molecule information must be applied. In the following text, when we refer to Linked-Read WGS (lrWGS) we are referring to the combination of biochemistry and algorithm approaches applied.

### Improvements in Linked-Read data

One limitation of the original GemCode approach was the need to combine the Linked-Read data with a standard short-read library for analysis. This was due to coverage imbalances seen in the GemCode library alone. To address this issue we modified the original biochemistry, replacing it with an isothermal amplification approach. The updated biochemistry now provides for more even genome coverage, approaching that of PCR-free short-read preparations (Figure 1).

**Figure 1:**
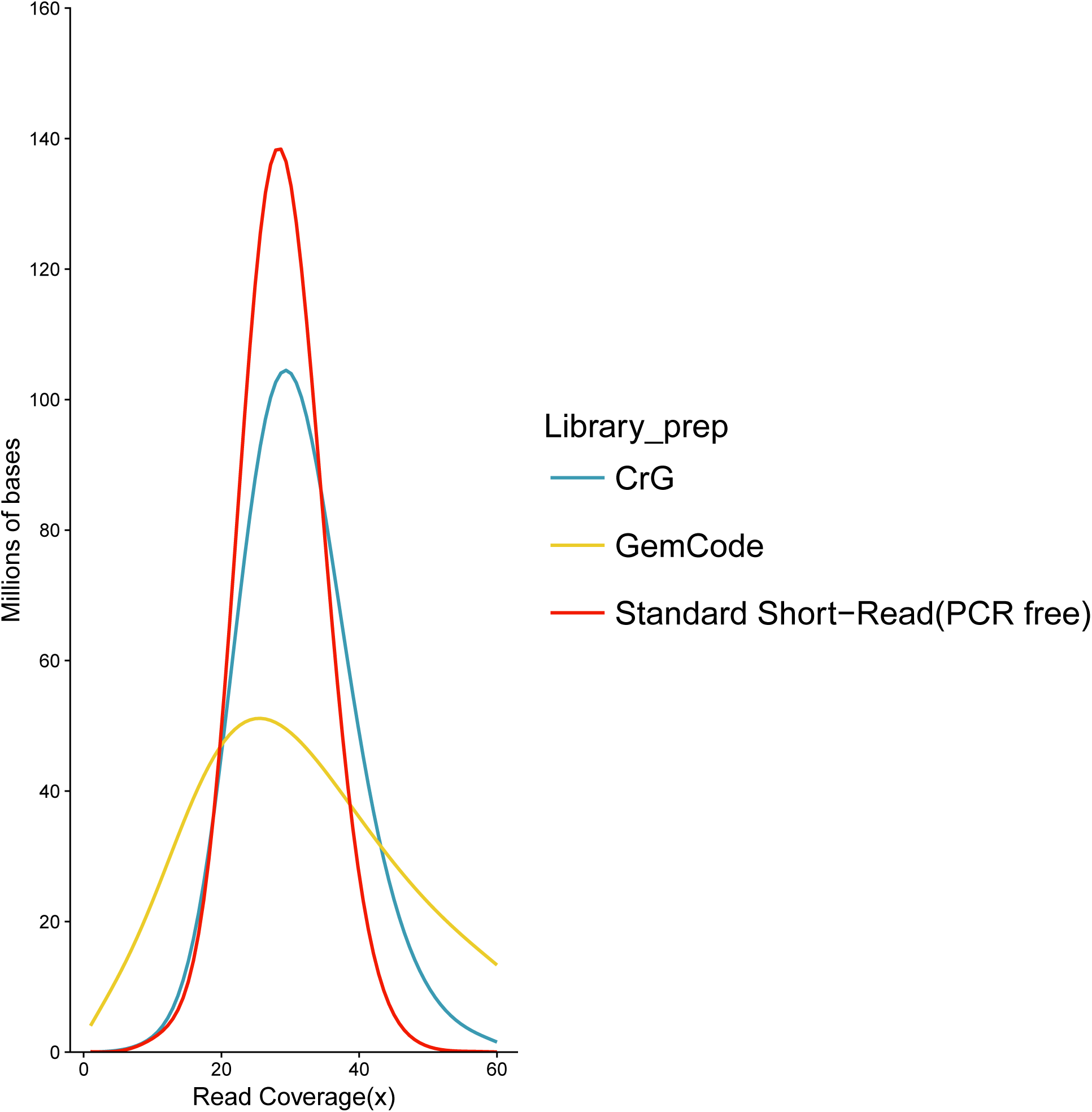
Coverage Evenness. Distribution of read coverage for the entire human genome (GRCh37). Comparisons between 10x Genomics Chromium Genome (CrG), 10x Genomics GemCode (GemCode), and Illumina TruSeq PCR-free standard short-read NGS library preparations (Standard Short Read (PCR-Free)). Sequencing was performed in an effort to match coverage (see methods). Note the shift of the CrG curve to the right, showing the improved coverage of Chromium vs. GemCode. X-axis represents the fold read coverage across the genome. Y-axis represents the total number of bases covered at any given read depth.

Additional improvements include increasing the number of barcodes from 737,000 to 4 million and the number of partitions from 100,000 to over 1 million. This allows for fewer DNA molecules per partition, or GEMs (Gelbead-in-EMulsion), and thus a greatly reduced background rate of barcode collisions: the rate at which two random loci occur in the same GEM (Supplemental Figure 1). The lowered background rate of barcode sharing increases the probability of correctly associating a short read to the correct molecule of origin, and increases the sensitivity for SV detection.

### Improved Genome and Exome Alignments

Several improvements were made in the Long Ranger analysis pipeline to better take advantage of the Linked-Read data type. The first of these, the Lariat™ aligner, expands on the ‘Read-Cloud’approach (Bishara et al. 2015). Lariat (https://github.com/10XGenomics/lariat) refines alignments produced by the BWA aligner by examining reads that map to multiple locations and determining if they share barcodes with reads that have high quality unique alignments (Li 2013). If a confident placement can be determined by taking advantage of the barcode information of the surrounding reads, the quality score of the correct alignment is adjusted (Supplemental Section 1). This approach allows for the recovery of 36-44 Mb of genome coverage when compared to PCR free short reads aligned following GATK best practices. Conversely, only 1-4 Mb of the genome has coverage in the PCR free data that is not seen using lrWGS (Figure 2). When looking at the genome distribution of these alignment gains, the amount of recovered alignments using lrWGS varies from chromosome to chromosome, but is consistent across samples (Supplemental Figure 2). This is due to genome structure, as the ability of lrWGS to rescue repetitive sequence, using the Lariat algorithm, depends on the repeats being far enough apart that they are not likely to share a barcode. Only in this case can the Lariat algorithm resolve reads mapping to multiple locations. The sequence gained using lrWGS is dominated by regions annotated as segmental duplication (roughly 75%), with the alignments to the decoy sequence accounting for another 13% and exonic sequences accounting for roughly 5% (Supplemental 1.2, Supplemental Table 1, Figure 2). Molecule length also impacts the amount of sequence recovered (Supplemental Figure 3).

**Table 1:** Summary of variant call numbers with respect to GIAB Table 1: The table shows the counts of variants (SNV and indel) from variant calls generated in four experiments: NA12878 Linked-Reads WGS data run through Long Ranger (NA12878 lrWGS), NA12878 TruSeq PCR-free data run through GATK-Best Practices pipeline (NA12878 srWGS), NA24385 Linked-Reads WGS data run through Long Ranger (NA24385 lrWGS), NA24385 TruSeq PCR-free data run through GATK-Best Practices pipeline (NA24385 srWGS). These variants were compared to the GIAB VCF truth set and GIAB BED confident regions using hap.py and data is shown per variant type for count of variants in the truth set and in the confident regions (along with sensitivity and specificity). Data is also shown for the same quantities when the variant calls were compared to the extended truth set (GIAB++ VCF) and the augmented confident region (GIAB++ BED).

|  | NA12878 lrWGS | NA12878 srWGS | NA24385 lrWGS | NA24385 srWGS |
| --- | --- | --- | --- | --- |
| Total Variants | 4,600,606 | 4,651,391 | 4,504,190 | 4,564,102 |
| Total SNVs | 3,808,856 | 3,760,296 | 3,731,448 | 3,689,866 |
| Sensitivity (SNVs) | 0.9965260 | 0.9978873 | 0.9972462 | 0.9984250 |
| Specificity (SNVs) | 0.9969829 | 0.9984747 | 0.9977549 | 0.9990125 |
| SNVs in confident regions | 3,153,057 | 3,152,799 | 3,053,304 | 3,053,249 |
| SNVs in truth set | 3,143,316 | 3,147,610 | 3,046,234 | 3,049,835 |
| Sensitivity (SNVs) (++) | 0.9944987 | 0.9954084 | 0.9966197 | 0.9973968 |
| Specificity (SNVs) (++) | 0.9745175 | 0.9879275 | 0.9703781 | 0.9838542 |
| SNVs in confident regions (++) | 3,266,048 | 3,224,849 | 3,151,491 | 3,111,146 |
| SNVs in truth set (++) | 3,182,558 | 3,185,469 | 3,057,434 | 3,059,818 |
| Total indels | 791,750 | 891,095 | 772,742 | 874,236 |
| Sensitivity (indels) | 0.9339752 | 0.9733969 | 0.9330855 | 0.9772879 |
| Specificity (indels) | 0.9501310 | 0.9820730 | 0.9493424 | 0.9851534 |
| Indels in confident regions | 361,547 | 368,216 | 347,786 | 354,897 |
| Indels in truth set | 334,577 | 348,699 | 321,517 | 336,748 |
| Sensitivity (indels) (++) | 0.9226400 | 0.9645790 | 0.9056345 | 0.9743154 |
| Specificity (indels) (++) | 0.9234368 | 0.9636761 | 0.8854908 | 0.9331947 |
| Indels in confident regions (++) | 379,399 | 383,935 | 474,879 | 491,054 |
| Indels in truth set (++) | 341,279 | 356,792 | 411,130 | 442,309 |

**Figure 2:**
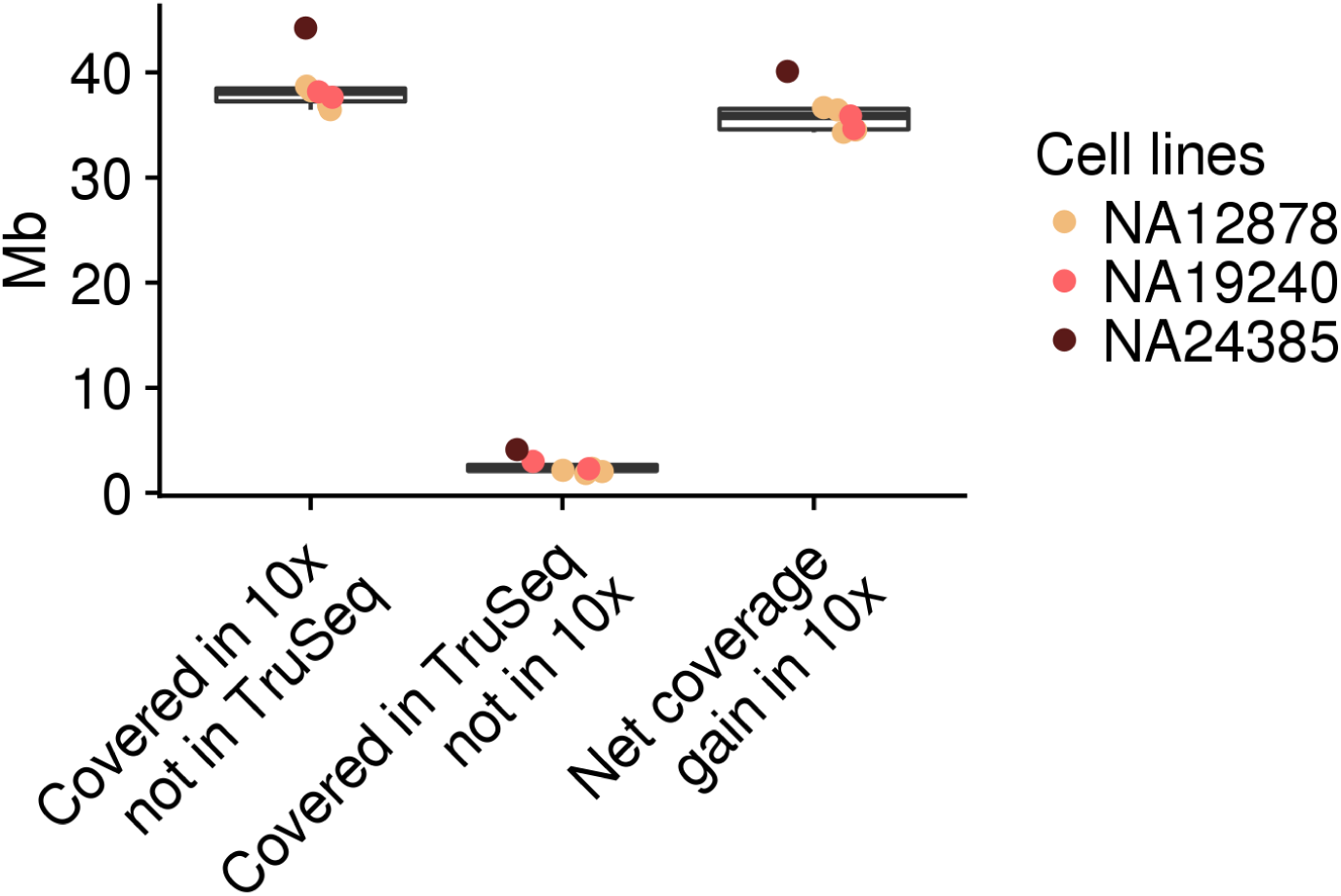
Comparison of unique genome coverage by assay. The y-axis shows the amount of sequence with a coverage of >=5 reads at MapQ >=30. Column 1 shows amount of the genome covered by 10x Chromium where PCR-free TruSeq does not meet that metric. Column 2 shows the amount of the genome covered by PCR-free TruSeq where 10x Chromium does not meet the metric. Column 3 shows the net gain of genome sequence with high quality alignments when using 10x Chromium versus PCR-free TruSeq. The comparison was performed on samples with matched sequence coverage (see methods).

When we look specifically at the ability of Lariat to improve read coverage over genes, we observe a net gain in gene coverage when performing lrWGS compared to srWGS, and even more robust improvement when performing lrWES compared to srWES (Supplemental Figure 4). When we limit the search space to a known set of 570 genes with closely related paralogs that confound short read alignment (NGS ‘dead zone’ genes (Mandelker et al. 2016)) we see a net gain in read coverage in 423 genes using lrWGS and 376 using lrWES. Further limiting the list to the 71 genes relevant to Mendelian disease, we see a net improvement in 51 of these genes using lrWGS and 41 genes using lrWES (Figure 3). Exome analysis was limited to multiple replicates of a single control sample, NA12878.

**Figure 3:**
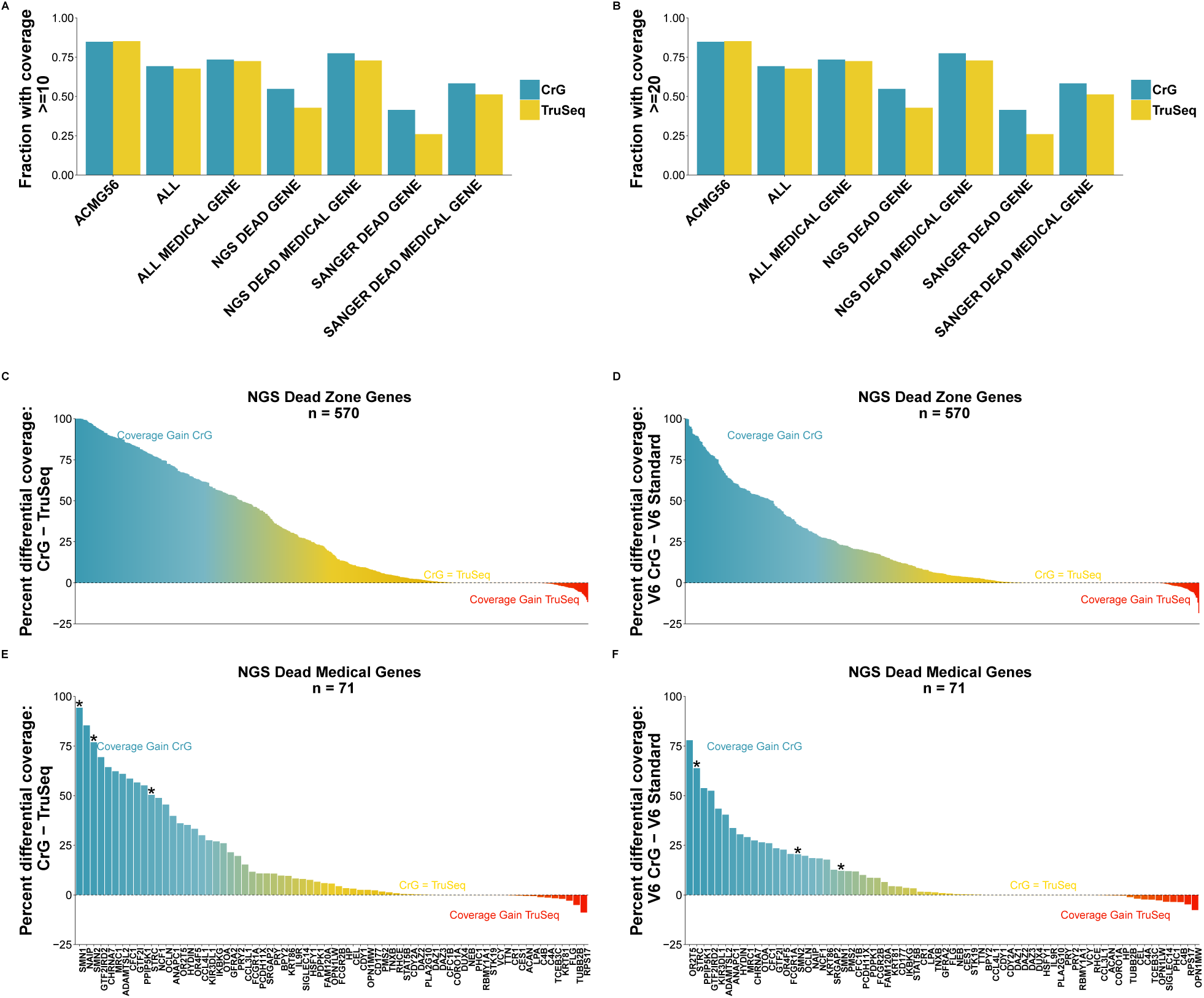
Gene finishing metrics. Gene finishing metrics for whole genome and whole exome sequencing across selected gene sets. Genome is shown on left, exome on right. Gene finishing is a metric used for expressing gene coverage and completeness. Finishing is defined as the percentage of exonic bases with at least 10x coverage for genome (Panel A) and at least 20x for exome (Panel B) (Mapping quality score >=MapQ30). CrG is Chromium Linked-Reads and TruSeq is PCR-free TruSeq. Top row: Gene finishing statistics for 7 disease relevant gene panels. Shown is the average value across all genes in each panel. While Chromium provides a coverage edge in all panel sets, the impact is particularly profound for ‘NGS Dead Zone’ genes. Panels C-F show the net coverage differences for individual genes when comparing Chromium to PCR-free TruSeq. Each bar shows the difference between the coverage in PCR-free TruSeq from the coverage in 10x Chromium. Panel C and D show the 570 NGS ‘dead zone’ genes for genome (panel C) and exome (panel D). Panels E and F limit the graphs to the list of NGS dead zone genes implicated in Mendelian disease. In panels C-F, the blue coloring highlights genes that are inaccessible to short read approaches, but accessible using CrG; the yellow coloring indicates genes where CrG is equivalent to short reads or provides only modest improvement. The red coloring shows genes with a slight coverage increase in TruSeq, though these genes are typically still accessible to CrG. Highlighted with an asterisk are the genes *SMN, SMN* and *STRC*. The comparison was performed on samples with matched coverage (see methods).

**Figure 4:**
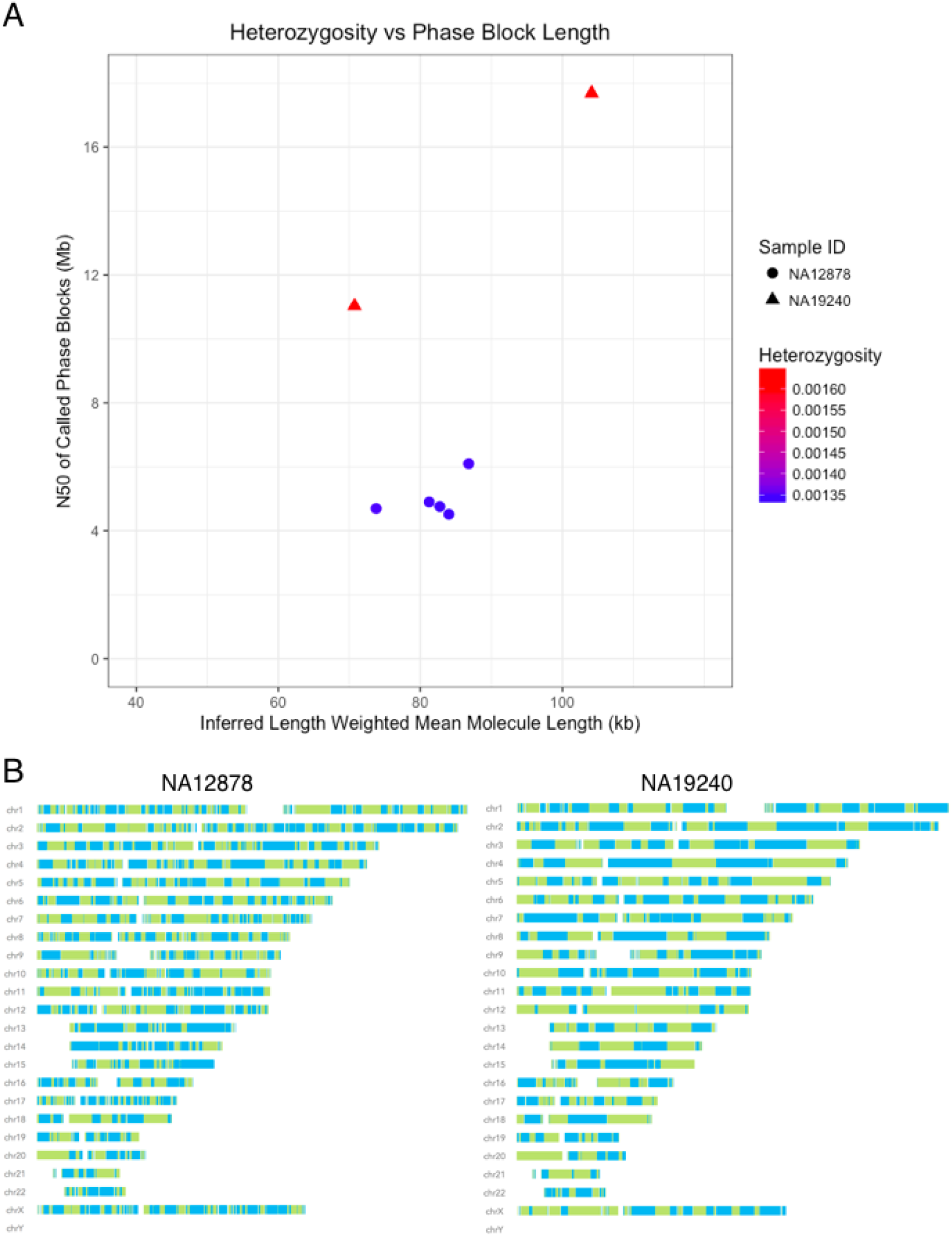
Haplotype reconstruction and phasing. A. Inferred Length weighted mean molecule length plotted against N50 of called Phase blocks (both metrics reported by Long Ranger) and differentiated by sample ID and heterozygosity. Heterozygosity was calculated by dividing the total number of heterozygous positions called by Long Ranger by the total number of non-N bases in the reference genome (GRCh37). Two replicates of NA19240 and 5 replicates of NA12878 were used. Samples with higher heterozygosity produce longer phase blocks than samples with less diversity when controlling for input molecule length. B. Phase block distributions across the genome for input length matched Chromium Genome samples NA12878 and NA19240. Phase blocks are shown as displayed in Loupe Genome Browser™. Solid colors indicate phase blocks.

### Small variant calling

Next, we assessed the performance of Linked-Reads for small variant calling (<50 bp). Small variant calling, particularly for single nucleotide variants (SNVs) outside of repetitive regions, is well powered by short reads because a high quality read alignment to the reference assembly is possible and the variant resides completely within the read. We used control samples, NA12878 and NA24385 as test cases. We produced two small variant call sets for each sample, one generated by running paired-end 10x Linked-Read Chromium libraries through the Long Ranger (lrWGS) pipeline and one produced by analyzing paired-end reads from a PCR-free TruSeq library using GATK pipeline (PCR-) following best practices recommendations: https://software.broadinstitute.org/gatk/best-practices/. We made a total of 4,585,361 PASS variant calls from the NA12878 lrWGS set, and 4,622,282 from the corresponding NA12878 srWGS set, with 4,436,102 calls in common to both sets (Table 1). Total numbers for both samples are in Table 1.

In order to assess the accuracy of the variant calling in each data set, we used the hap.py tool (Krusche)(https://github.com/Illumina/hap.py, commit 6c907ce) to compare the lrWGS and PCR-VCFs to the Genome in a Bottle (GIAB) high confidence call set (v. 3.2.2) (Zook et al. 2014). We chose this earlier version as it was the last GIAB data set that did not include 10x data as an input for their call set curation. This necessitated the use of GRCh37 as a reference assembly rather than the more current GRCh38 reference assembly. This limited us to analyzing only the 82.67% of SNV calls that overlap the high confidence regions. Initial results suggested that the lrWGS calls had comparable sensitivity (>99.65%) and specificity (>99.70%) for SNVs (Table 1). We observed slightly diminished indel sensitivity (>93.31%) and specificity (>94.93%), driven largely by regions with extreme GC content and low complexity sequences (LCRs). Recent work suggests indel calling is still a challenging problem for many approaches, but that only 0.5% of LCRs overlap regions of the genome thought to be functional based on annotation or conservation (Li et al. 2017). Additionally, we compared the sensitivity of homozygous and heterozygous calls (Supplemental Table 2). Both lrWGS and PCR-have higher sensitivity and specificity for homozygous alternate variants than heterozygous variants.

**Table 2:** Gene, variant distance and RVIS score for clinically-relevant genes Table 2: Impact of molecule length and constraint on the ability of Linked-Reads to phase causative variants. As molecule length increases within a sample, the likelihood that two causative variants will be phased relative to each other also increases. However, genes that are not highly constrained (e.g. *TTN*) are more likely to show phasing between distant variants at small molecule lengths because more heterozygous variants are likely to occur between those variants than in highly constrained genes.

| Sample | Gene | Var1 | Var2 | Var distance | RVIS score | RVIS % | Molecule length | Var phased? |
| --- | --- | --- | --- | --- | --- | --- | --- | --- |
| B12-38 | DYSF | chr2:71,778,243dupT | chr2:71,817,342_71,817,343delinsAA | 39,097 bp | -1.31 | 4.65% | 13,553 bp | No |
| B12-38 | DYSF | chr2:71,778,243dupT | chr2:71,817,342_71,817,343delinsAA | 39,097 bp | -1.31 | 4.65% | 16,911 bp | No |
| B12-38 | DYSF | chr2:71,778,243dupT | chr2:71,817,342_71,817,343delinsAA | 39,097 bp | -1.31 | 4.65% | 18,439 bp | No |
| B12-38 | DYSF | chr2:71,778,243dupT | chr2:71,817,342_71,817,343delinsAA | 39,097 bp | -1.31 | 4.65% | 18,461 bp | Yes |
| B12-38 | DYSF | chr2:71,778,243dupT | chr2:71,817,342_71,817,343delinsAA | 39,097 bp | -1.31 | 4.65% | 19,309 bp | Yes |
| B12-38 | DYSF | chr2:71,778,243dupT | chr2:71,817,342_71,817,343delinsAA | 39,097 bp | -1.31 | 4.65% | 21,226 bp | Yes |
| B12-38 | DYSF | chr2:71,778,243dupT | chr2:71,817,342_71,817,343delinsAA | 39,097 bp | -1.31 | 4.65% | 34,800 bp | Yes |
| B12-38 | DYSF | chr2:71,778,243dupT | chr2:71,817,342_71,817,343delinsAA | 39,097 bp | -1.31 | 4.65% | 42,939 bp | Yes |
| B12-38 | DYSF | chr2:71,778,243dupT | chr2:71,817,342_71,817,343delinsAA | 39,097 bp | -1.31 | 4.65% | 85,077 bp | Yes |
| B12-38 | DYSF | chr2:71,778,243dupT | chr2:71,817,342_71,817,343delinsAA | 39,097 bp | -1.31 | 4.65% | 88,410 bp | Yes |
| B12-38 | DYSF | chr2:71,778,243dupT | chr2:71,817,342_71,817,343delinsAA | 39,097 bp | -1.31 | 4.65% | 119,747 bp | Yes |
| B12-38 | DYSF | chr2:71,778,243dupT | chr2:71,817,342_71,817,343delinsAA | 39,097 bp | -1.31 | 4.65% | 130,101 bp | Yes |
| B12-112 | POMT2 | chr14:77,745,107A>G | chr14:77,778,305C>T | 33,198 bp | -0.93 | 9.68% | 10,609 bp | No |
| B12-112 | POMT2 | chr14:77,745,107A>G | chr14:77,778,305C>T | 33,198 bp | -0.93 | 9.68% | 12277 bp | No |
| B12-112 | POMT2 | chr14:77,745,107A>G | chr14:77,778,305C>T | 33,198 bp | -0.93 | 9.68% | 15,536 bp | No |
| B12-112 | POMT2 | chr14:77,745,107A>G | chr14:77,778,305C>T | 33,198 bp | -0.93 | 9.68% | 16,546 bp | No |

Table 2: Gene, variant distance and RVIS score for clinically-relevant genes (*continued*)
| Sample | Gene | Var1 | Var2 | Var distance | RVIS score | RVIS % | Molecule length | Var phased? |
| --- | --- | --- | --- | --- | --- | --- | --- | --- |
| B12-112 | POMT2 | chr14:77,745,107A>G | chr14:77,778,305C>T | 33,198 bp | -0.93 | 9.68% | 20,782 bp | No |
| B12-112 | POMT2 | chr14:77,745,107A>G | chr14:77,778,305C>T | 33,198 bp | -0.93 | 9.68% | 21,106 bp | No |
| B12-112 | POMT2 | chr14:77,745,107A>G | chr14:77,778,305C>T | 33,198 bp | -0.93 | 9.68% | 21,858 bp | No |
| B12-112 | POMT2 | chr14:77,745,107A>G | chr14:77,778,305C>T | 33,198 bp | -0.93 | 9.68% | 54,569 bp | Yes |
| B12-112 | POMT2 | chr14:77,745,107A>G | chr14:77,778,305C>T | 33,198 bp | -0.93 | 9.68% | 55,546 bp | Yes |
| B12-112 | POMT2 | chr14:77,745,107A>G | chr14:77,778,305C>T | 33,198 bp | -0.93 | 9.68% | 107,082 bp | Yes |
| B12-112 | POMT2 | chr14:77,745,107A>G | chr14:77,778,305C>T | 33,198 bp | -0.93 | 9.68% | 112,692 bp | Yes |
| B12-21 | TTN | chr2:179,585,773C>A | chr2:179,531,966C>A | 53,807 bp | 2.17 | 98.04% | 17,432 bp | Yes |
| B12-21 | TTN | chr2:179,585,773C>A | chr2:179,531,966C>A | 53,807 bp | 2.17 | 98.04% | 18,128 bp | Yes |
| B12-21 | TTN | chr2:179,585,773C>A | chr2:179,531,966C>A | 53,807 bp | 2.17 | 98.04% | 18,158 bp | Yes |
| B12-21 | TTN | chr2:179,585,773C>A | chr2:179,531,966C>A | 53,807 bp | 2.17 | 98.04% | 20,756 bp | Yes |
| B12-21 | TTN | chr2:179,585,773C>A | chr2:179,531,966C>A | 53,807 bp | 2.17 | 98.04% | 28,799 bp | Yes |
| B12-21 | TTN | chr2:179,585,773C>A | chr2:179,531,966C>A | 53,807 bp | 2.17 | 98.04% | 29,796 bp | Yes |
| B12-21 | TTN | chr2:179,585,773C>A | chr2:179,531,966C>A | 53,807 bp | 2.17 | 98.04% | 47,443 bp | Yes |
| B12-21 | TTN | chr2:179,585,773C>A | chr2:179,531,966C>A | 53,807 bp | 2.17 | 98.04% | 63,218 bp | Yes |
| B12-21 | TTN | chr2:179,585,773C>A | chr2:179,531,966C>A | 53,807 bp | 2.17 | 98.04% | 64,199 bp | Yes |
| B12-21 | TTN | chr2:179,585,773C>A | chr2:179,531,966C>A | 53,807 bp | 2.17 | 98.04% | 67,034 bp | Yes |

Table 2: Impact of molecule length and constraint on the ability of Linked-Reads to phase causative variants.
| Sample | Gene | Var1 | Var2 | Var distance | RVIS score | RVIS % | Molecule length | Var phased? |
| --- | --- | --- | --- | --- | --- | --- | --- | --- |
| B12-21 | TTN | chr2:179,585,773C>A | chr2:179,531,966C>A | 53,807 bp | 2.17 | 98.04% | 90,767 bp | Yes |
| B12-21 | TTN | chr2:179,585,773C>A | chr2:179,531,966C>A | 53,807 bp | 2.17 | 98.04% | 93,253 bp | Yes |
| UC-394 | TTN | chr2:179,584,098C>T | chr2:179,395,221T>A | 188,877 bp | 2.17 | 98.04% | 13,118 bp | Yes |
| UC-394 | TTN | chr2:179,584,098C>T | chr2:179,395,221T>A | 188,877 bp | 2.17 | 98.04% | 16,791 bp | No |
| UC-394 | TTN | chr2:179,584,098C>T | chr2:179,395,221T>A | 188,877 bp | 2.17 | 98.04% | 18,192 bp | No |
| UC-394 | TTN | chr2:179,584,098C>T | chr2:179,395,221T>A | 188,877 bp | 2.17 | 98.04% | 18,841 bp | No |
| UC-394 | TTN | chr2:179,584,098C>T | chr2:179,395,221T>A | 188,877 bp | 2.17 | 98.04% | 28,033 bp | No |
| UC-394 | TTN | chr2:179,584,098C>T | chr2:179,395,221T>A | 188,877 bp | 2.17 | 98.04% | 30,653 bp | No |
| UC-394 | TTN | chr2:179,584,098C>T | chr2:179,395,221T>A | 188,877 bp | 2.17 | 98.04% | 32,530 bp | No |
| UC-394 | TTN | chr2:179,584,098C>T | chr2:179,395,221T>A | 188,877 bp | 2.17 | 98.04% | 69,939 bp | Yes |
| UC-394 | TTN | chr2:179,584,098C>T | chr2:179,395,221T>A | 188,877 bp | 2.17 | 98.04% | 87,045 bp | Yes |
| UC-394 | TTN | chr2:179,584,098C>T | chr2:179,395,221T>A | 188,877 bp | 2.17 | 98.04% | 88,605 bp | Yes |
| UC-394 | TTN | chr2:179,584,098C>T | chr2:179,395,221T>A | 188,877 bp | 2.17 | 98.04% | 89,863 bp | Yes |

The GIAB high confidence data set is known to be quite conservative and we wished to explore whether there was evidence for variants called outside of the GIAB set in the lrWGS. We utilized publicly available 40x coverage PacBio data sets available from the GIAB consortium (Zook et al. 2016) to evaluate Linked-Read putative false positive variant calls. Initial manual inspection of 25 random locations suggested that roughly half of the hap.py identified lrWGS false positive calls were well supported by short read or PacBio evidence, and were haplotype consistent in lrWGS and were likely called false positive due to deficiencies in the GIAB truth set (Supplemental Table 3). We then did a global analysis of all 9,513 SNV and 18,030 indel putative false positive calls identified in NA12878 and looked for evidence of the alternate alleles in aligned PacBio reads only. This analysis provided evidence that 2,377 SNV and 12,812 indels of the GIAB determined false positive calls were likely valid calls (Supplemental Figure 5, Supplemental File 1). This prompted us to extend our analysis to include 69.72 Mb for NA12878 and 70.66 Mb for NA24385 of the genome corresponding to regions of 2-6-fold degeneracy as determined by the ‘CRG Alignability track’ in addition to the GIAB defined confident regions (see Methods for details on GIAB++ BED). We reanalyzed the variant calls with the hap.py tool with the augmented confident regions. This allowed us to identify an additional 19,688 SNV and 5,444 indels as true positives. We anticipate that this is a conservative estimate since our hap.py defined false positive calls are inflated due to little or no PacBio or short-read coverage in many of these regions. Of the total putative false positive calls exclusive to the GIAB++ analysis, 61.95% (45,665) of SNVs and 42.08% (4,637) of indels could not be validated because of little or no PacBio read coverage (Supplemental Figure 5). These data show the lrWGS approach provides for the identification of more small variants than can be identified by short read only approaches, driven by an increase in the percentage of the genome for which lrWGS can obtain high quality alignments (see Table 1).

**Table 3:** SV Intersections Table 3: Different intersections of Long Ranger SV calls with a ground truth dataset published (Parikh et al. 2016). Comparison class identified in the leftmost column. Large deletions (>=30kb) intersected against all deletions >=30kb in the ground truth set. Smaller deletions (<30kb), marked as PASS by our algorithm, intersected against the full deletion ground truth set.

|  | Query<br>Number | Query<br>Overlap | Target<br>Number | Target<br>Overlap |
| --- | --- | --- | --- | --- |
| $\geq 30\text{kb}$ | 17 | 8 | 11 | 8 |
| $< 30\text{kb}$ | 5136 | 2024 | 2294 | 2024 |

**Figure 5:**
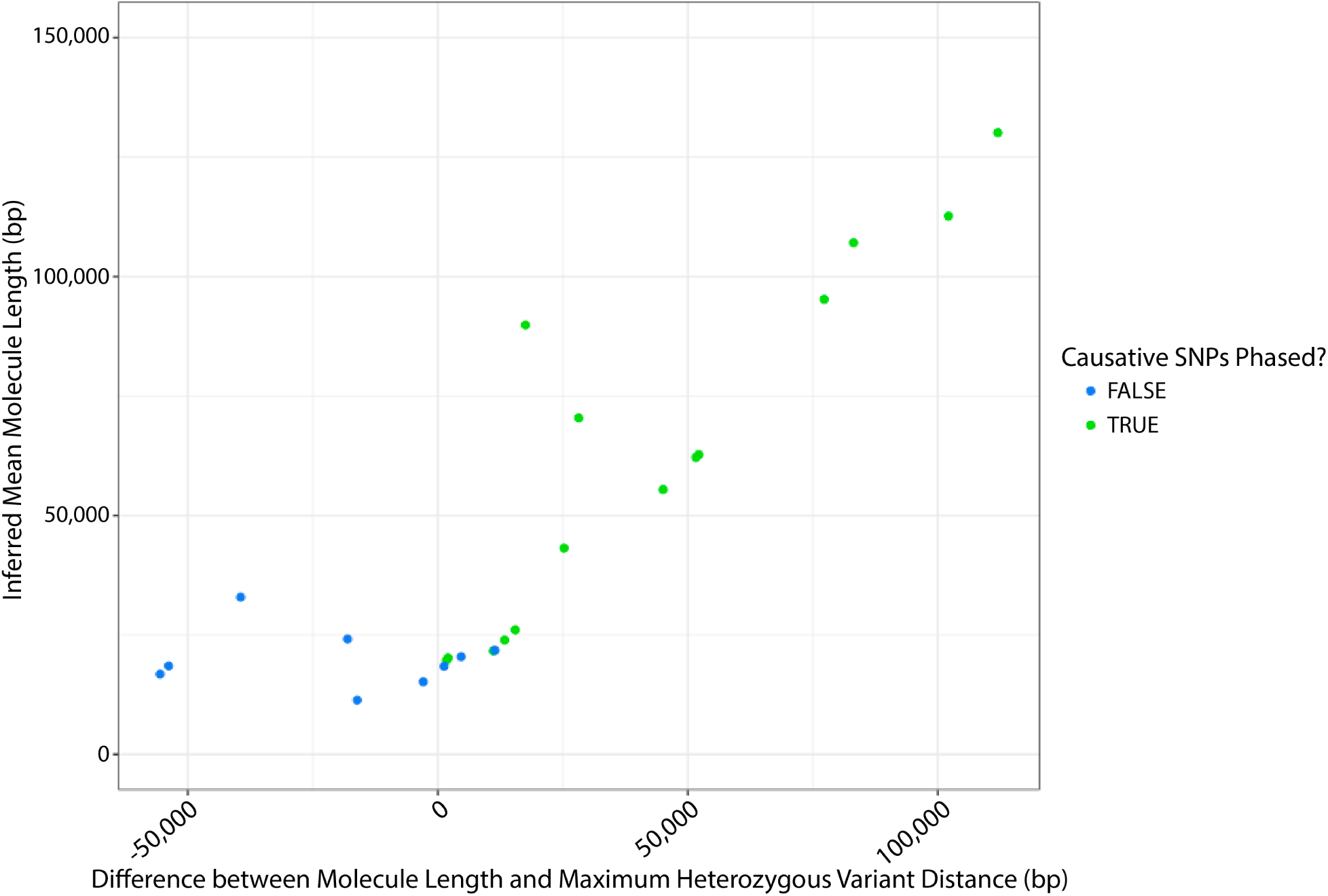
Validated examples of impact of molecule length on phasing (7.25Gb). Blue dots represent samples for which the variants of interest are not phased, and green dots represent samples for which there is phasing of the variants of interest. At longer molecule lengths (>50kb), the molecule length was always longer than the maximum distance between heterozygous SNPs in a gene, and phasing between the causative SNPs was always observed. As molecule length shortens, it becomes more likely that the maximum distance between SNPs exceeds the molecule length (reflected as a negative difference value) and phasing between the causative SNPs was never observed in these cases. When maximum distance is similar to the molecule length causative SNPs may or may not be phased. This is likely impacted by the molecule length distribution within the sample.

### Haplotype reconstruction and phasing

An advantage of Linked-Reads is the ability to reconstruct multi-megabase haplotypes from genome sequence data (called phase blocks) for a single sample. Haplotype reconstruction increases sensitivity for calling heterozygous variants, particularly SVs (Huddleston et al. 2016). It also improves variant interpretation by providing information on the physical relationship of variants, such as whether variants within the same gene are in *cis* or *trans*. In the control samples analyzed, we see phase block N50 values for lrWGS of 10.3 Mb for NA12878, 9.58Mb for NA24385, 16.8 Mb for NA19240 and 302 kb for lrWES using Agilent SureSelect v6 baits on NA12878. This allowed for complete phasing of 91.1% for NA12878 genome, 90.9% for NA24385 genome, and 91.0% for NA19240 genome, and an average of 91% for NA12878 exome. Phase block length is a function of input molecule length, molecule size distribution and of sample heterozygosity extent and distribution. At equivalent mean molecule lengths, phase blocks will be longer in more diverse samples (Figure 4, Supplemental Figure 6). For samples with similar heterozygosity, longer input molecules will increase phase block lengths (Supplemental Figure 7).

**Figure 6:**
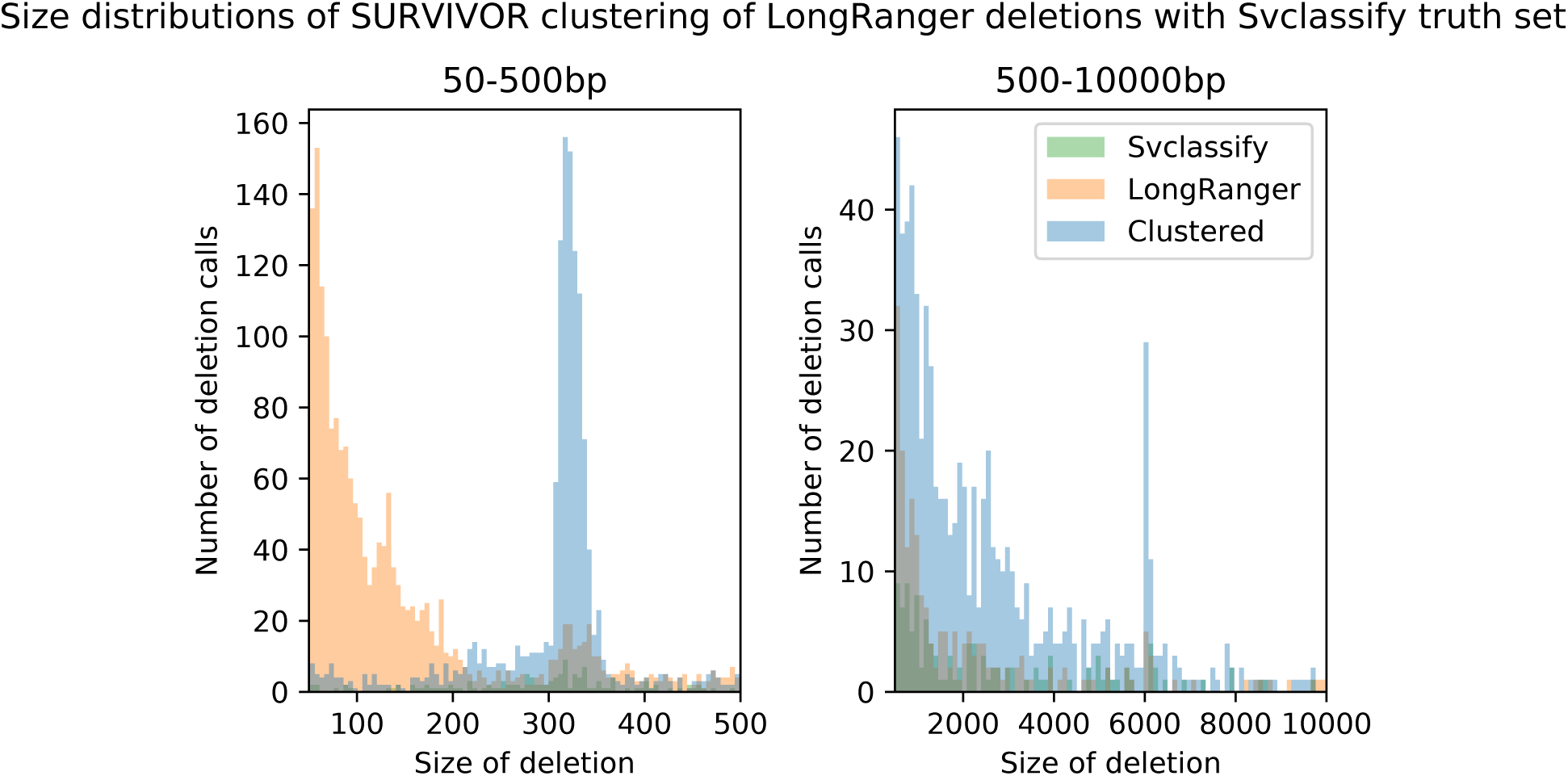
Deletions size distributions Long Ranger calls intersected with the svclassify truth set by size. True positive calls are blue, false negative calls are green and false positive calls are orange. Most false positives are present in the <250bp size range, reflecting the lack of small deletions in the svclassify set. Peaks corresponding to Alu and L1/L2 elements can be seen at ∼320bp and ∼6kbp respectively.

We assessed the accuracy of our phasing calls by comparing the Linked-Read phasing results to published phasing results derived from pedigree sequencing. We compare our NA12878 results with the Illumina Platinum genomes (Eberle et al. 2017) phasing results derived from jointly phasing the 17 member CEPH pedigree. Following the previous analysis (Amini et al. 2014), we decompose phasing errors into “short-switches” and “long-switches”. Short-switches are defined by a small number of isolated variants incorrectly phased, whereas “long-switches” are those errors in which an incorrect junction is formed that persists for many variants across a longer distance. The rate of each switch type is measured per phased heterozygous variant. We also measure 1) the rate at which a given SNP is correctly phased to other variants in its phase block (which heavily penalizes long switch errors inside large phase blocks), and 2) for SNPs inside a gene boundary, the rate at which a SNP inside a gene is correctly phased to other variants in the gene. Independent studies have demonstrated that Linked-Read phasing has best in class accuracy compared to a variety of other phasing methods (Chaisson et al. 2017; Choi et al. 2018). Short switch error rates average ∼0.0002, long switch error rates average ∼2e-5, and within-phase-block correct rate has an average of ∼0.98. See Supplemental Table 4 for details.

**Table 4:**
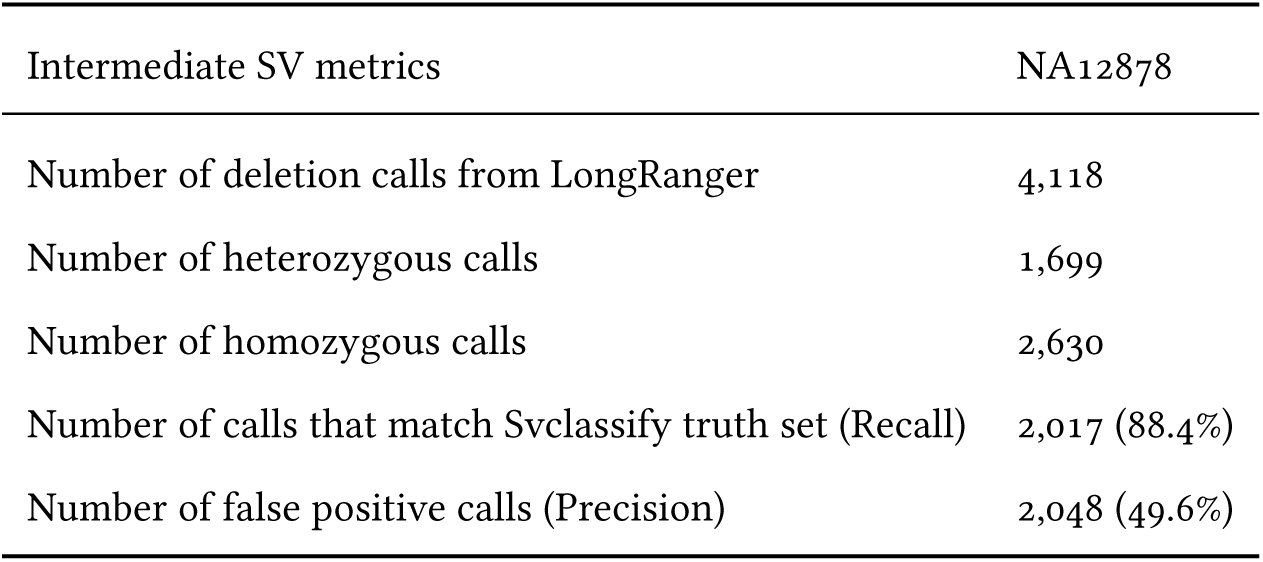
Intermediate SV Calls Table 4: Intermediate SV (50bp to 30kbp) results. The number of calls generated by the intermediate SV algorithms are reported and broken down by inferred zygosity. SURVIVOR (Jeffares et al. 2017) was used to merge these intermediate SVs with the svclassify (Parikh et al. 2016) truth set which had also been subsetted to the same size range, and the resulting true positive and false positive rates are reported as well as the associated recall and precision.

| Intermediate SV metrics | NA12878 |
| --- | --- |
| Number of deletion calls from LongRanger | 4,118 |
| Number of heterozygous calls | 1,699 |
| Number of homozygous calls | 2,630 |
| Number of calls that match Svclassify truth set (Recall) | 2,017 (88.4%) |
| Number of false positive calls (Precision) | 2,048 (49.6%) |

Phase block construction using lrWES is additionally constrained by the bait set used to perform the capture and the reduced variation seen in coding sequences. In order to analyze factors impacting phase block construction, we assessed four samples with known compound heterozygous variants in three genes known to cause Mendelian disease, *DYSF, POMT*_*2*_, and *TTN*. The variant separation ranged from 33 Kb to over 188 Kb (Table 2). Initial DNA extractions yielded long molecules ranging in mean size from 75 Kb - 112 Kb. We analyzed these samples using the Agilent SureSelect V6 exome bait set, with downsampling of sequence data to both 7.25 Gb (∼60x coverage) and 12 Gb of sequence (∼100x coverage). In all cases, the variants were phased with respect to each other and determined to be in *trans*, as previously determined by orthogonal assays. By comparing the phasing of NA12878 Linked-Read exome data to phasing determined from pedigree analysis of the Illumina Platinum Genomes CEPH pedigree (including NA12878) we are able to determine that the global probability a SNP is phased correctly within a gene ranges from 99.95-99.99%, making mis-phasing of two heterozygous variants in a gene relative to each other a very rare event.

In three of the four cases, the entire gene was phased. The *DYSF* gene was not completely phased in any sample because the distance between heterozygous SNPs at the 3’ end of the gene was substantially longer than the mean molecule length. This gene is in the top 5% of genes intolerant to variation as determined by the RVIS metric, a measure of evolutionary constraint, suggesting that reduced exonic heterozygosity over the gene would be a common occurrence impairing complete phasing (Petrovski et al. 2013).

Many samples of interest have already been extracted using standard methods not optimized for high molecular weight DNA and may not be available for a fresh re-extraction to obtain DNA optimized for length. For this reason, we wanted to understand the impact of reduced molecule length on our ability to phase the genes and variants in these samples. We took the original freshly extracted long molecules and sheared them to various sizes, aiming to assess lengths ranging from 5Kb to the original full length samples (Table 2). These results illustrate the complex interplay between molecule length distribution and the observed heterozygosity within a region. For example, in sample B12-21, with variants in *TTN* that are 53 Kb apart, the variants could be phased, even with the smallest molecule size. However in sample B12-122, with variants in *POMT*_*2*_ only 33 Kb apart, variant phasing is lost at 20 Kb DNA lengths. This appeared to be due to a higher rate of heterozygous variation in *TTN* allowing the phasing of distant heterozygous sites to occur by phasing the many other heterozygous variants that occurred between them. A general lack of variation in *POMT*_*2*_ precluded such stitching together of shorter molecules by phasing of intermediate heterozygous variation. To confirm this, we assessed the maximum distance between heterozygous sites observed in each gene. We then plotted the difference between the inferred molecule length and this distance and against the molecule length and assessed the impact on causative SNP phasing (Figure 5). In general, when the maximum distance between heterozygous SNPs is greater than the molecule length (negative values), the ability to phase causative SNPs decreases. There are exceptions to this as the longer molecules in the molecule size distribution will sometimes allow tiling between the variants, therefore extending phase block size beyond what would be expected based on the mean length alone.

Linked-Reads allow for the reconstruction of long haplotypes, or phase blocks. Optimizing for long input molecules provides for maximum phase block size, but even shorter molecule lengths can provide gene level phasing. Further, in the context of sequencing for the identification of disease, causative heterozygous variants would be expected to aid in the phasing of the disease-causing gene as they would provide the necessary variation to distinguish the two haplotypes.

### Structural variant detection

Structural variants remain one of the most difficult types of variation to accurately ascertain, in part because they tend to cluster in duplicated and repetitive regions, but also because the various signals for these events can be challenging to detect with short reads. Accurate and specific SV detection is challenging due in large part to the limitations of assessing long range information using short reads, which only provide information over short distances. Another complicating factor is the many types of structural variants, each requiring the detection of a different signal depending on the type and mechanism of the event (Alkan et al. 2011; Collins et al. 2017). There is increasing evidence that grouping reads by their source haplotype improves SV sensitivity, but this is not commonly done in practice (Huddleston et al. 2016; Chaisson et al. 2017). It is of interest to identify the full range of SVs, particularly larger SVs as these larger events are more frequently associated with changes in gene expression signatures (Chiang et al. 2017).

#### Large-scale SVs (>30K)

Long Ranger uses two novel algorithms to identify large SVs, one that assesses deviations from expected barcode coverage and one that looks for unexpected barcode overlap between distant regions. The barcode coverage algorithm is useful for assessing CNVs, while the barcode overlap method can detect a variety of SVs, but fails to detect terminal events (See Supplemental Section 3). SV calls are a standard output of the Long Ranger pipeline and are described using standard file formats. We used two approaches to assess lrWGS performance on large SVs. First, we compared SV calls from the NA12878 sample to validated calls described in a recent publication of a structural variant classifier, svclassify (Parikh et al. 2016). Next, we obtained the GeT-RM CNVPanel, a collection of known events including large deletions, duplications, inversions, balanced translocations and unbalanced translocations designed to assess performance of clinical aCGH.

Long Ranger identifies event types by matching to simple models of deletions, duplications and inversions. Therefore, there are additional events where Long Ranger identifies clear evidence for anomalous barcode overlap, but is unable to match the event to one of the pre-defined models. These undefined events are rendered as unknown and represent deficiencies in SV annotation. The validated call set published with svclassify (Parikh et al. 2016) contains deletions and insertions, but no balanced events. By contrast, the Long Ranger pipeline output contains deletions, duplications and balanced events, but Long Ranger does not currently call insertions (Supplemental Table 5).

**Table 5:** Gene, variant type and pipeline call for clinically-relevant genes Table 5: Ability of Linked-Reads to call variation in samples with known exon-level deletions and duplications. Exome or whole genome sequencing was used on samples freshly extracted from cell lines or on archival DNA samples. The ability of the barcode-aware algorithms to call exon-level events is completely dependent on phasing. Longer DNA length and increased sequencing coverage sometimes improve variant calling, but this appears to be rescued by enabling phasing.

| Sample | Gene | Variant type | Source | Assay | Calc mean<br>length | Region<br>phased? | Called by >=1<br>method? |
| --- | --- | --- | --- | --- | --- | --- | --- |
| A | MSH2 | Single Exon Duplication | Archival DNA | SureSelectV6, 7.25Gb (60x) | 64kb | No | No |
| A | MSH2 | Single Exon Duplication | Archival DNA | SureSelectV6, 12Gb (100x) | 53kb | No | No |
| B | PMS2 | Single Exon Duplication | Archival DNA | SureSelectV6, 7.25Gb (60x) | 65kb | Yes | Yes |
| B | PMS2 | Single Exon Duplication | Archival DNA | SureSelectV6, 12Gb (100x) | 59kb | Yes | Yes |
| C | BRCA1 | Single Exon Duplication | Cell line | SureSelectV6, 7.25Gb (60x) | 96kb | No | No |
| C | BRCA1 | Single Exon Duplication | Cell line | SureSelectV6, 12Gb (100x) | 78kb | No | No |
| C | BRCA1 | Single Exon Duplication | Cell line | Whole Genome, 128Gb (30x) | 88kb | No | No |
| C | BRCA1 | Single Exon Duplication | Archival DNA | SureSelectV6, 7.25Gb (60x) | 28kb | No | No |
| C | BRCA1 | Single Exon Duplication | Archival DNA | SureSelectV6, 12Gb (100x) | 27kb | No | No |
| D | BRCA2 | Single Exon Duplication | Archival DNA | SureSelectV6, 7.25Gb (60x) | 24kb | No | No |
| D | BRCA2 | Single Exon Duplication | Archival DNA | SureSelectV6, 12Gb (100x) | 19kb | Yes | Yes |
| E | BRCA1 | Two exon deletion | Cell line | SureSelectV6, 7.25Gb (60x) | 106kb | No | No |
| E | BRCA1 | Two exon deletion | Cell line | SureSelectV6, 12Gb (100x) | 98kb | No | No |
| E | BRCA1 | Two exon deletion | Archival DNA | SureSelectV6, 7.25Gb (60x) | 71kb | No | No |
| E | BRCA1 | Two exon deletion | Archival DNA | SureSelectV6, 12Gb (100x) | 80kb | No | No |
| F | BRCA1 | Two exon deletion | Cell line | SureSelectV6, 7.25Gb (60x) | 97kb | Yes | Yes |
| F | BRCA1 | Two exon deletion | Cell line | SureSelectV6, 12Gb (100x) | 107kb | Yes | Yes |

Table 5: Ability of Linked-Reads to call variation in samples with known exon-level deletions and 667 duplications.
| Sample | Gene | Variant type | Source | Assay | Calc mean<br>length | Region<br>phased? | Called by $\geq 1$<br>method? |
| --- | --- | --- | --- | --- | --- | --- | --- |
| F | BRCA1 | Two exon deletion | Archival DNA | SureSelectV6, 7.25Gb (60x) | 15kb | No | No |
| F | BRCA1 | Two exon deletion | Archival DNA | SureSelectV6, 12Gb (100x) | 12kb | Yes | Yes |
| G | PMS2 | Two exon deletion | Archival DNA | SureSelectV6, 7.25Gb (60x) | 57kb | Yes | Yes |
| G | PMS2 | Two exon deletion | Archival DNA | SureSelectV6, 12Gb (100x) | 48kb | Yes | Yes |
| H | PMS2 | 2-3 exon deletion | Archival DNA | SureSelectV6, 7.25Gb (60x) | 54kb | Yes | Yes |
| H | PMS2 | 2-3 exon deletion | Archival DNA | SureSelectV6, 12Gb (100x) | 42kb | Yes | Yes |
| I | PMS2 | Large structural variant | Archival DNA | SureSelectV6, 7.25Gb (60x) | 43kb | Yes | No |
| I | PMS2 | Large structural variant | Archival DNA | SureSelectV6, 12Gb (100x) | 35kb | Yes | No |
| I | PMS2 | Large structural variant | Archival DNA | Whole genome, 128Gb (30x) | 28kb | Yes | No |
| I | MSH2 | Two exon deletion | Archival DNA | SureSelectV6, 7.25Gb (60x) | 43kb | No | No |
| I | MSH2 | Two exon deletion | Archival DNA | SureSelectV6, 12Gb (100x) | 35kb | No | No |
| I | MSH2 | Two exon deletion | Archival DNA | Whole genome, 128Gb (30x) | 28kb | Yes | Yes |

We first consider deletion variants >30 Kb. There are 11 of these in the svclassify set and 17 in the Long Ranger PASS set, with 8 being common to both (Table 3). All of the variants that match svclassify events also show Mendelian consistency and breakpoint agreement within the CEU/CEPH trio. Of the three svclassify calls not called by Long Ranger, one is called by Long Ranger as an event <30kb, one is called but filtered to the candidate list due to overlap with a segmental duplication, and one is an error in the svclassify set relative to GRCh37.p13 (Supplemental Section 4.1). We checked for Mendelian consistency in the 9 events unique to the Long Ranger set. Eight of these events showed consistent inheritance, though two had inconsistent breakpoints when compared to the parents (Supplemental Table 6). One of these breakpoint inconsistent events entirely contains a breakpoint consistent event on the same haplotype. The second breakpoint inconsistent event overlaps an additional inheritance-consistent Long Ranger call, and thus represents a failure of the algorithm to annotate the event as being a more complex event. The final event called by Long Ranger, but not showing inheritance consistency, is a call in the telomeric region of chr2 that overlaps a known reference assembly issue. The call appears to be made due to a drop in phased coverage on one haplotype immediately adjacent to a known reference gap, and is likely a false positive.

We next tested 23 samples with 29 validated balanced or unbalanced SVs from the GeT-RM CNVPanel available from Coriell. These cell lines have multiple, orthogonal assays confirming the presence of their described structural variants. We detected 27 of the 29 structural variants, correctly characterizing 22 of the 23 samples tested (Supplemental Table 7). One additional event was in the ‘candidate’ SV list as it overlaps a segmental duplication, which are known problematic regions for SV calling. The missed event is a balanced translocation with a breakpoint in a heterochromatic region of chromosome 16. This region is represented by Ns in the reference assembly and will be invisible to any sequence-based method relying on the reference genome (Schneider et al. 2017).

We also assessed the impact of sequence depth on large SV identification. Deletion and duplication signals were detectable with as little as 5Gb (∼1x genomic read coverage) (Supplemental Figure 8). Balanced events required roughly 50Gb of sequence for the algorithm to call these events, though signal in the data suggested algorithmic improvements could lessen this requirement (Supplemental Figure 9).

#### Intermediate SV Calls (50bp - 30Kb)

We next considered deletions between 50 bp and 30 Kb in the NA12878 sample. The Long Ranger pipeline was run using GATK and thus we can obtain two sets of files: deletion and insertion calls from GATK that are approximately 250bp or less, and deletion calls from Long Ranger algorithms. As Long Ranger only calls deletions, we only considered these calls in the following analysis. We also ran the LUMPY (Layer et al., 2014) algorithm using the developer recommendations but without tuning parameters (Supplemental Table 8: SuppTable8_IntSVs). We obtained 1,824 deletion calls from GATK and 4,118 from Long Ranger, with 1,699 of these being heterozygous (Table 4). This compares to 6,965 deletions >50bp per sample in a study combining the output of 13 different algorithms on short read data (Chaisson et al. 2017). This same study also used long reads to identify 9,488 deletions >50bp per sample, underscoring the challenges of identifying these events with short reads.

Using only the output of Long Ranger, we compared our calls to the calls in svclassify. We identified 2,017 calls (88.4%), with 2,048 (49.6%) labeled as false positives (Table 4). Combining the GATK and Long Ranger calls keeps recall roughly the same, but lowers the precision roughly 10% (Supplemental Table 8). Of note, the Long Ranger calls provide improved detection of larger SVs, with an expected bump around 300 bp, likely accounted for by better representation of ALUs (Figure 6).

While sensitivity of the Long Ranger approach is good, this comes at the expense of specificity (Table 4, Supplemental Table 8). Given the bias in specificity in phased versus unphased regions, we expect that algorithmic improvements will produce further gains in sensitivity and specificity for this class of variants. Additionally, we suspect the small number of events <200 bp in the svclassify set is not representative of the true number of calls in a given sample.

Linked-Reads provide improvements for SV detection over standard short read approaches. However, there is ample room for algorithmic improvement using SVs. For example, approaches based on local reassembly could be utilized for insertion discovery.

### Analysis of samples from individuals with inherited disease

We went on to investigate the utility of Linked-Read analysis on samples with known variants. In particular, we were interested in events that are typically difficult with a standard, short read exome. We were able to obtain samples from a cohort that had been assessed using a high depth NGS-based inherited predisposition to cancer screening panel. This cohort contained samples with known exon level deletion and duplication events. We analyzed these 12 samples from 9 individuals using an Agilent SureSelect V6 Linked-Read exome at both 7.25 Gb (equivalent to ∼60x raw coverage) and 12 Gb (∼100x) coverage (Table 5). For three samples patient-derived cell lines were available in addition to archival DNA, allowing us to investigate the impact of DNA length on exon-level deletion/duplication calling.

We were able to identify 5 of the 9 expected exon-level events in these samples in at least one sample type/depth combination. In 2 samples, increasing depth to 12Gb enabled calling that was not possible at 7.25Gb (Samples D and F (archival), Table 5). For the three samples with matched cell lines and archival DNA, two had variants that could not be called in either sample type at either depth, while sample F could be called at both depths for the longer DNA extracted from the cell line, but could only be called at the higher depth in the shorter archival sample. Because the algorithms for calling these variants are written to make use of phasing and barcode information, there is a striking correlation between the ability to phase the gene and to call the variant, with no variants successfully called in samples that could not be phased over the region of interest.

For two of the samples where Linked-Read exome sequencing was unable to phase or call the known variant, we performed lrWGS. In one case, the presence of intronic heterozygous variation was able to restore phasing to the gene and the known event was called. In the second case, there was still insufficient heterozygous variation in the sample to allow phasing and the event was not called. This again demonstrates that phasing is dependent both on molecule length as well as sample heterozygosity. Some samples in this group had decreased diversity in the regions of interest compared to other samples, and we were less likely to be able to call variants in these samples. (Supplemental Figure 10). Generally, it should be possible to increase the probability of phasing a gene in an exome assay by augmenting the bait set to provide coverage for very common (MAF > 25%) intronic variant SNPs, thus preserving the cost savings of exome analysis, but increasing the power of the Linked-Reads to phase. The number of additional probes could be minimized with long molecules. Despite this, samples with generally reduced heterozygosity will remain difficult to phase and completely characterize. However, the addition of read coverage-based algorithms, such as those used with standard short read exome sequencing, would likely increase sensitivity in unphased regions.

One sample in this set contained both a single exon event and a large variant in the *PMS2* gene. Despite phasing the *PMS2* gene we were unable to call this variant in either genome or exome sequencing. Manual inspection of the data reveals increased phased barcode coverage in the *PMS*_*2*_ region, supporting the presence of a large duplication that was missed by the SV calling algorithms (Supplemental Figure 11). This indicates room for additional improvements in the variant calling algorithms.

Linked-Reads provide a better first line approach than standard short read assays to assess individuals for variants in these genes. While we were not able to identify 100% of the events, we were able to identify 5 of 9 of these events using a standard exome based approach, rather than a specialized assay. Improved baiting approaches, the addition of standard short read algorithms, or WGS should improve that ability to identify these variants. Lastly, there is room for algorithmic improvement as at least one variant had clear signal in the Linked-Read data, but failed to be recognized by current algorithms.

## Discussion

Short read sequencing has become the workhorse of human genomics. This cost effective, high throughput, and accurate base calling approach provides robust analysis of short variants in unique regions of the genome, but struggles to reliably call SVs, cannot assess variation across the entire genome, and fails to reconstruct long range haplotypes (Sudmant et al. 2015). Recent studies have highlighted the importance of including haplotype information and more complete SV identification in genome studies (Chaisson et al. 2017, 2017). Analyzing human genomes in their diploid context will be a critical step forward in genome analysis (Aleman 2017). Toolkits that support the representation of sequence and variation, a necessary component of supporting true, diploid assembly, are now becoming available (Garrison et al. 2018). We have described an improved implementation of Linked-Reads, a method that improves the utility of short read sequencing. The increased number of partitions and improved biochemistry mean a single Linked-Read library, constructed from ∼1 ng of DNA, can be used for genome analysis. This approach, coupled with novel algorithms in Long Ranger, allows short reads to reconstruct multi-megabase phase blocks, identify large balanced and unbalanced structural variants, and identify small variants, even in regions of the genome typically recalcitrant to short read approaches.

Some limitations to this approach currently exist. We observe a loss of coverage in regions of the genome that show extreme GC content. We additionally see reduced performance in small indel calling, though this largely occurs in homopolymer regions and LCRs. Recent work suggests ambiguity in such regions may be tolerated for a large number of applications (Li et al. 2017). Although Linked-Reads can resolve many repetitive elements and genome regions, highly repetitive sequences that are larger than the length of input DNA are not resolvable by Linked-Reads. This limitation is common to all technologies currently available, including long-read sequencing. Repeat copies that reside on the same molecule will be subject to the same limitations as standard short read approaches. It is also clear that algorithmic improvements to Long Ranger would improve variant calling, particularly as some classes of variants, such as insertions, are not yet attempted. However, this is not uncommon for new data types and there has already been some progress in this area (Spies et al. 2016; Elyanow et al. 2017; Xia et al. 2017; Karaoglanoglu et al. 2018). An additional limitation in this study is the reliance on a reference sample for calling variants, which creates reference bias and the inability to call variants in regions that are not resolved in the reference, as was the case with the structural variant in the pericentric region on chromosome 16. To bypass any reference bias, Linked-Read data can also be used to perform diploid *de novo* assembly in combination with an assembly program, Supernova (Weisenfeld et al. 2017).

Despite these limitations, Linked-Read sequencing provides a clear advantage over short reads alone. This pipeline allows for the construction of long range haplotypes as well as the identification of short variants and SVs from a single library and analysis pipeline. No other approach, to our knowledge, that scales to thousands of genomes provides this level of detail for genome analysis. Other recent studies have demonstrated the power of Linked-Reads to resolve complex variants in both germline and cancer samples (Collins et al. 2017; Greer et al. 2017; Viswanathan et al.; Nordlund et al. 2018). Recent work demonstrates that Linked-Reads outperforms the switch accuracy and phasing completeness of other haplotyping methods, and provides multi-MB phase blocks (Chaisson et al. 2017). In another report, Linked-Reads and the Supernova assembly algorithm have been used to perform de novo assembly on 17 individuals to identify novel sequence (Wong et al. 2018). The ability to provide reference free analysis promises to increase our understanding of diverse populations. Finally, the ability to represent and analyze genomes in terms of haplotypes, rather than compressed haploid representations, represents a crucial shift in our approach to genomics, allowing for a more complete and accurate reconstruction of individual genomes.

## Methods

*Samples and DNA Isolation* Control samples (NA12878, NA19240, NA24385, NA19240, and NA24385) were obtained as fresh cultured cells from the Coriell Cell biorepository (https://catalog.coriell.org/1/NIGMS). DNA was isolated using the Qiagen MagAttract HMW DNA kit and quantified on a Qubit fluorometer following recommended protocols: https://support.10xgenomics.com/genome-exome/index/doc/user-guide-chromium-genome-reagent-kit-v2-chemistry.

Samples with known large SVs were obtained as cell lines from the NIGMS Human Genetic Cell Repository at the Coriell Institute for Medical Research (repository ID numbers are listed in Table s1). Frozen cell pellets were thawed rapidly at 37°C in 1mL PBS. High molecular weight DNA was then extracted following recommended protocols, as above.

Clinical samples from individuals with known heterozygous variants in three Mendelian disease loci (*DYSF, POMT2* and *TTN*) were collected at the Massachusetts General Hospital, Analytic and Translational Genetics Unit and shipped to 10x genomics as cell lines. Genomic DNA was extracted from each cell line as described above. Use of samples from the Broad Institute was approved by the Partners IRB (protocol 2013P001477).

Clinical samples from individuals with inherited cancer were collected at The Institute of Cancer Research, London and shipped to 10x genomics as cell lines or archival DNA. This sample cohort was previously accessed for predisposition to cancer. Samples were recruited through the Breast and Ovarian Cancer Susceptibility (BOCS) study and the Royal Marsden Hospital Cancer Series (RMHCS) study, which aimed to discover and characterize disease predisposition genes. All patients gave informed consent for use of their DNA in genetic research. The studies have been approved by the London Multicentre Research Ethics Committee (MREC/01/2/18) and Royal Marsden Research Ethics Committee (CCR1552), respectively. Samples were also obtained through clinical testing by the TGLclinical laboratory, an ISO 15189 accredited genetic testing laboratory. The consent given from patients tested through TGLclinical includes the option of consenting to the use of samples/data in research; all patients whose data was included in this study approved this option. DNA was extracted from cell lines as described above and archival DNA samples were checked for size and quality according to manufacturer’s recommendations: https://support.10xgenomics.com/genome-exome/sample-prep/doc/demonstrated-protocol-hmw-dna-qc.

*Chromium™ Linked-Read Library Preparation* 1.25 ng of high molecular weight DNA was loaded onto a Chromium controller chip, along with 10x Chromium reagents (either v1.0 or v2.0) and gel beads following recommended protocols: https://assets.contentful.com/an68im79xiti/4z5JA3C67KOyCE2ucacCM6/d05ce5fa3dc4282f3da5ae7296f2645b/CG00022_GenomeReagentKitUserGuide_RevC.pdf. The initial part of the library construction takes place within droplets containing beads with unique barcodes (called GEMs). The library construction incorporates a unique barcode that is adjacent to read one. All molecules within a GEM get tagged with the same barcode, but because of the limiting dilution of the genome (roughly 300 haploid genome equivalents) the chances that two molecules from the same region of the genome are partitioned in the same GEM is very small. Thus, the barcodes can be used to statistically associate short reads with their source long molecule.

Target enrichment for the Linked-Read whole exome libraries was performed using Agilent Sure Select V6 exome baits following recommended protocols: https://assets.contentful.com/an68im79xiti/Zm2u8VlFa8qGYW4SGKG6e/4bddcc3cd60201388f7b82d241547086/CG000059_DemonstratedProtocolExome_RevC.pdf. Supplemental Figure 12 describes targeted sequencing with Linked-Reads.

### GemCode™ Linked-Read Library Preparation

For the GemCode comparator analyses, Linked-Read libraries were prepared for truth samples NA12878, NA12877, and NA12882 using a GemCode controller and GemCode V1 reagents following published protocols (Zheng et al. 2016).

### TruSeq PCR-free Library Preparation

350-800 ng of genomic DNA was sheared to a size of ∼385 bp using a Covaris^®^M220 Focused Ultrasonicator using the following shearing parameters: Duty factor = 20%, cycles per burst = 200, time = 90 seconds, Peak power 50. Fragmented DNA was then cleaned up with 0.8x SPRI beads and left bound to the beads. Then, using the KAPA Library Preparation Kit reagents (KAPA Biosystems, Catalog # KK8223), DNA fragments bound to the SPRI beads were subjected to end repair, A-base tailing and Illumina^®^‘PCR-free’ TruSeq adapter ligation (1.5 *µ*M final concentration of adapter was used). Following adapter ligation, two consecutive SPRI cleanup steps (1.0X and 0.7X) were performed to remove adapter dimers and library fragments below ∼150 bp in size. No library PCR amplification enrichment was performed. Libraries were then eluted off the SPRI beads in 25 ul elution buffer and quantified with quantitative PCR using KAPA Library Quant kit (KAPA Biosystems, Catalog # KK4824) and an Agilent Bioanalyzer High Sensitivity Chip (Agilent Technologies) following the manufacturer’s recommendations.

Target enrichment for the Linked-Read whole exome libraries was performed using Agilent Sure Select V6 exome baits following recommended protocols.

*Sequencing* Libraries were sequenced on a combination of Illumina^®^instruments (HiSeq^®^2500, HiSeq 4000, and HiSeq X). Paired-End sequencing read lengths were as follows: TruSeq and Chromium whole genome libraries (2X150bp); Chromium whole exome libraries (2X100bp or 114bp, 98bp), and Gemcode libraries (2X98bp). lrWGS libraries are typically sequenced to 128 Gb, compared to 100 Gb for standard TruSeq PCR-free libraries. The additional sequence volume compensates for sequencing the barcodes as well a small number of additional sources of wasted data and gives an average, de-duplicated coverage of approximately 30x. To demonstrate the extra sequence volume is not the driver of the improved alignment coverage, we performed a gene finishing comparison at matched volume (100Gb lrWGS and 100Gb TruSeq PCR-) and continue to see coverage gains (Supplemental Figure 12).

## Analysis

*Comparison of 10X and GATK Best Practices* We ran the GATK Best practices pipeline to generate variant calls for TruSeq PCR-free data using the latest GATK3.8 available at the time. We first subsample the reads to obtain 30x whole genome coverage. The read set is then aligned to GRCh37, specifically the hg19-2.2.0 reference using BWA-MEM (version 0.7.12). The reads are then sorted, the duplicates are marked, and the bam is indexed using picard tools (version 2.9.2). We then perform indel realignment and recalibrate the bam (base quality score recalibration) using known indels from Mills Gold Standard and 1000G project and variants from dbsnp (version 138). Finally we call both indel and SNVs from the bam using HaplotypeCaller and genotype it to produce a single vcf file. This vcf file is then compared using hap.py (https://github.com/Illumina/hap.py, commit 6c907ce) to the truth variant set curated by Genome in a Bottle on confident regions of the genome. We calculate sensitivity and specificity for both SNVs and indels to contrast the fidelity of the Long Ranger short variant caller and the GATK-Best Practices pipeline. All Long Ranger runs were performed with a pre-release build of Long Ranger version 2.2 utilizing GATK as a base variant caller. Long Ranger 2.2 adds a large-scale CNV caller that employs barcode coverage information and incremental algorithmic improvements. Long Ranger 2.2 has since been released.

### Development of extended truth set

Any putative false positive variant found in the TruSeq/GATK or Chromium/Long Ranger VCFs, was tested for support in the PacBio data. Raw PacBio FASTQs were aligned to the reference using BWA-MEM-x pacbio (Li 2013). To test a variant, we fetch all PacBio reads covering the variant position, and retain the substring aligned within 50bp of the variant on the reference. We re-align the PacBio read sequence to the +/-50bp interval of the reference, and the same interval with the alternate allele applied. A read is considered to support the alternate allele if the alignment score to the alt-edited template exceeds the alignment score of the reference template. A variant was considered to be validated if at least 2 PacBio reads supported the alt allele, at least 10 PacBio reads covered the locus, and the overall alternate allele fraction seen in the PacBio reads was at least 25%.

We selected regions of 2-6 fold degeneracy as determined by the ‘CRG Alignability’ track (Derrien et al. 2012) as regions where improved alignment is likely to yield credible novel variants. We took the union of the GIAB confident regions BED file with these regions to determine the GIAB++ confident regions BED. The amount of sequence added to the GIAB++ BED differs by sample, as the original GIAB confident regions are sample specific.

### *Structural variant comparison against deletion ground truth* After segmenting the Long Ranger deletion calls by size, we overlapped them to the svclassify set (Parikh et al. 2016) using the bedr package and bedtools v2.27.1 (Quinlan and Hall 2010). We retained for further analysis those

>30kb showing at least 50% reciprocal overlap. We also searched for Mendelian inheritance patterns on NA12878’s parents (NA12891 and NA12892) in these large SVs and breakpoint co-location. We annotated 8 overlapping events and they showed almost perfect breakpoint and Mendelian inheritance agreement within the CEU/CEPH trio. All their genotypes were phased too. In the svclassify overlapping deletions, all of the breakpoints except for the 3’ most in chr5:104,432,114-104,503,672 had a read’s length distance from each other. We then curated the remaining 9 events called by Long Ranger that were not in the svclassify set. Of notice is that one event (chr1:189,704,517-189,783,347) is contained within a larger deletion (chr1:189,690,000-189,790,000). Among the non-overlapping deletions, were six large SVs presenting breakpoint and Mendelian consistency in the phased genotypes. The other three (chr1:189,690,000-189,790,000; chr11:55,360,000-55,490,000; chr2:242,900,000-243,080,000) had very different breakpoints, unphased but consistent genotypes or no support from the parents.

We took the Long Ranger deletion calls between 50bp and 30kb generated by both Long Ranger algorithms and GATK and merged them using SURVIVOR (Jeffares et al. 2017) allowing variants up to 50bp apart to be merged. SURVIVOR was used again with a 50bp merge distance to merge the Long Ranger deletion callset with deletions in the svclassify set. The resulting merged VCFs were then parsed to determine overlap and support for Long Ranger calls.

## Acknowledgements

We thank the individuals who donated specimens for research (EGA Accession EGAD00001004319). This manuscript would not have been possible without their contributions. We thank Stephane Boutet and Sarah Taylor for reviewing the manuscript. We also wish to thank Kariena Dill for help with manuscript preparation, Kevin Wu for assistance with the technical aspects on setting up markdown and Docker. We also thank Jamie Schwendinger-Schreck for project management as well as invaluable contributions in manuscript preparation. All read data sequenced under this study is deposited at the Short Read Archive under accession number PRJNA428496. Genomic short variation and structural variant study data is deposited at the European Variation Archive under accession PRJEB28297.

